# The microtubule plus-end tracking protein Bik1 is required for chromosome congression

**DOI:** 10.1101/2021.06.16.447861

**Authors:** Alexander Julner, Marjan Abbasi, Victoria Menéndez-Benito

## Abstract

During mitosis, sister chromatids congress on both sides of the spindle equator to facilitate the correct partitioning of the genomic material. Chromosome congression requires a finely tuned control of microtubule dynamics by the kinesin motor proteins. In *Saccharomyces cerevisiae*, the kinesin proteins Cin8, Kip1, and Kip3 have a pivotal role in chromosome congression. It has been hypothesized that additional proteins that modulate microtubule dynamics are also involved. Here, we show that the microtubule plus-end tracking protein Bik1 – the budding yeast ortholog of CLIP-170 – is essential for chromosome congression. We find that nuclear Bik1 localizes to the kinetochores in a cell-cycle-dependent manner. Disrupting the nuclear pool of Bik1 with a nuclear export signal (Bik1-NES) leads to a slower cell cycle progression characterized by a delayed metaphase-anaphase transition. Bik1-NES cells have mispositioned kinetochores along the spindle in metaphase. Furthermore, using proximity-dependent methods, we identify Cin8 as an interaction partner of Bik1. Deleting CIN8 reduces the amount of Bik1 at the spindle. In contrast, Cin8 retains its typical bilobed distribution in the Bik1-NES mutant and does not localize to the unclustered kinetochores. Thus, we propose that Bik1 functions with Cin8 to regulate kinetochore-microtubule dynamics for correct kinetochore positioning and chromosome congression.

## Introduction

Cell division relies on the accurate segregation of sister chromatids to generate functional daughter cells. Before segregation, sister chromatids are positioned at the spindle equator through a process known as chromosome congression that culminates with the metaphase plate formation in metazoans (reviewed in (Maiato et al., 2017)). The establishment of a metaphase plate forces all the chromosomes to segregate from the same starting position, thereby contributing to faithful segregation. Despite the relevance of chromosome congression, its molecular mechanisms remain unclear.

The budding yeast *Saccharomyces cerevisiae* is an appealing model organism to study spindle dynamics because it is amenable to genetic engineering and has a relatively simple spindle. Budding yeast has 16 chromosomes. After duplication, each sister chromatid is linked by a kinetochore to a single kinetochore microtubule emanating from the spindle pole body (SPB), the yeast centrosome (Winey & O’Toole, 2001). Each SPB also gives rise to a few interpolar microtubules (starting with around six and ending with two interpolar microtubules from each SPB), with plus-ends that overlap at the middle of the spindle (O’Toole et al., 1999; Peterson & Ris, 1976; Winey et al., 1995). In prometaphase, kinetochores congress in two opposite clusters near the spindle equator, equidistant from the SPBs. As in metazoan cells, this appears to occur prior to biorientation of sister chromatids (Marco et al., 2013).

Several studies have shown that kinetochore microtubules are regulated in a length-dependent manner (Gardner et al., 2005; Pearson et al., 2006; Sprague et al., 2003). Short microtubules tend to grow, whereas long microtubules that extend towards the spindle equator are destabilized and depolymerize. As sister chromatids are held together by cohesin, depolymerization of kinetochore microtubules generates tension, mediated by the Ndc80 and Dam1 complexes’ ability to track the dynamic tips of the kinetochore microtubules (Powers *et al*., 2009; Suzuki *et al*., 2016).

Kinesin motors – proteins that move along microtubules powered by the hydrolysis of adenosine triphosphate (ATP) – are involved in chromosome congression in several organisms (see Maiato *et al*., 2017, Table 1). In budding yeast, the deletion of the kinesins Cin8, Kip3, and, to a lesser extent, Kip1, results in declustering of kinetochores (Gardner et al., 2008; Marco et al., 2013; Tytell & Sorger, 2006; Wargacki et al., 2010). Cin8 and Kip3 have been shown to act as length-dependent microtubule depolymerases, promoting catastrophe of long microtubules (Gardner et al., 2008; Su et al., 2011; Varga et al., 2006).

**Table 1.** A subset of the protein interactors of Bik1-FLAG (See Data S1 for a complete list).

| Name | % Coverage | Unique peptides |
| --- | --- | --- |
| STU2 | 72.33 | 104 |
| BIM1 | 67.00 | 21 |
| CIN8 | 50.67 | 46 |
| KIP2 | 49.33 | 28 |
| KAR9 | 47.33 | 26 |
| KIP1 | 38.67 | 28 |
| KAR3 | 38.67 | 23 |
| CIK1 | 22.67 | 10 |
| VIK1 | 21.00 | 10 |
| KIP3 | 15.33 | 6 |

It has long been speculated that other microtubule-associated proteins could play a role in chromosome congression (Kops *et al*., 2010). Specifically, microtubule plus-end tracking proteins (+TIPs) are interesting candidates, because they accumulate at kinetochore microtubule plus-ends and regulate their dynamics (reviewed in Akhmanova & Steinmetz, 2010). Bik1, the yeast ortholog of CLIP-170, is a +TIP protein that exists in cytoplasmic and nuclear pools within the cell (Carvalho et al., 2004). The cytoplasmic pool is important for mitotic spindle positioning through both the Kar9 and the dynein pathways (reviewed in Miller *et al*., 2006). First, Bik1 promotes phosphorylation of Kar9 for asymmetric localization at the old SPB (Moore et al., 2006). Kar9, together with the +TIP protein Bim1 and the actin-guided myosin motor protein Myo2, mediates the capture of astral microtubules at the bud cortex, contributing to spindle alignment and nuclear migration (Xiang, 2018). During anaphase, Bik1 is transported by the kinesin Kip2 to the plus-ends of astral microtubules, where it facilitates the recruitment of dynein (Carvalho *et al*., 2004; Roberts *et al*., 2014). Dynein is a minus end-directed motor and will pull the spindle resulting in nuclear migration to the bud.

The nuclear pool of Bik1 is associated with the spindle at SPBs and kinetochores (Carvalho et al., 2004; Lin et al., 2001), but its function is not well understood. Early studies found that deleting BIK1 results in cells with short or non-detectable astral microtubules (Berlin *et al*., 1990). In contrast, cells overexpressing Bik1 display short spindle microtubules and long astral microtubules. In this study, we investigate the nuclear function of Bik1 and its role in chromosome congression and cell cycle progression.

## Results and discussion

### Bik1 localizes to kinetochores in a cell cycle-dependent manner

To better understand the nuclear pool of Bik1, we first explored the localization of the protein at different stages of the cell cycle. We used cells expressing Bik1-superfolder GFP (sfGFP), mTurquoise-Tub1 (alpha tubulin, microtubule marker), and either Ndc80-tagRFPT (kinetochore marker) or Spc42-tagRFPT (SPB marker) (Figure 1A). In metaphase cells, nuclear Bik1 was primarily found between SPBs and overlapping with kinetochores. This bilobed kinetochore-associated distribution has also been reported for the kinesins Cin8, Kip1, and Kip3 (Gardner et al., 2008; Tytell & Sorger, 2006; Wargacki et al., 2010). Anaphase cells, on the other hand, displayed nuclear Bik1 concentrated in patches along interpolar microtubules, again similar to what is reported for kinesins Cin8, Kip1, and Kip3. Cytoplasmic Bik1, meanwhile, was localized to astral microtubule tips at the cell cortex.

**Figure 1.**
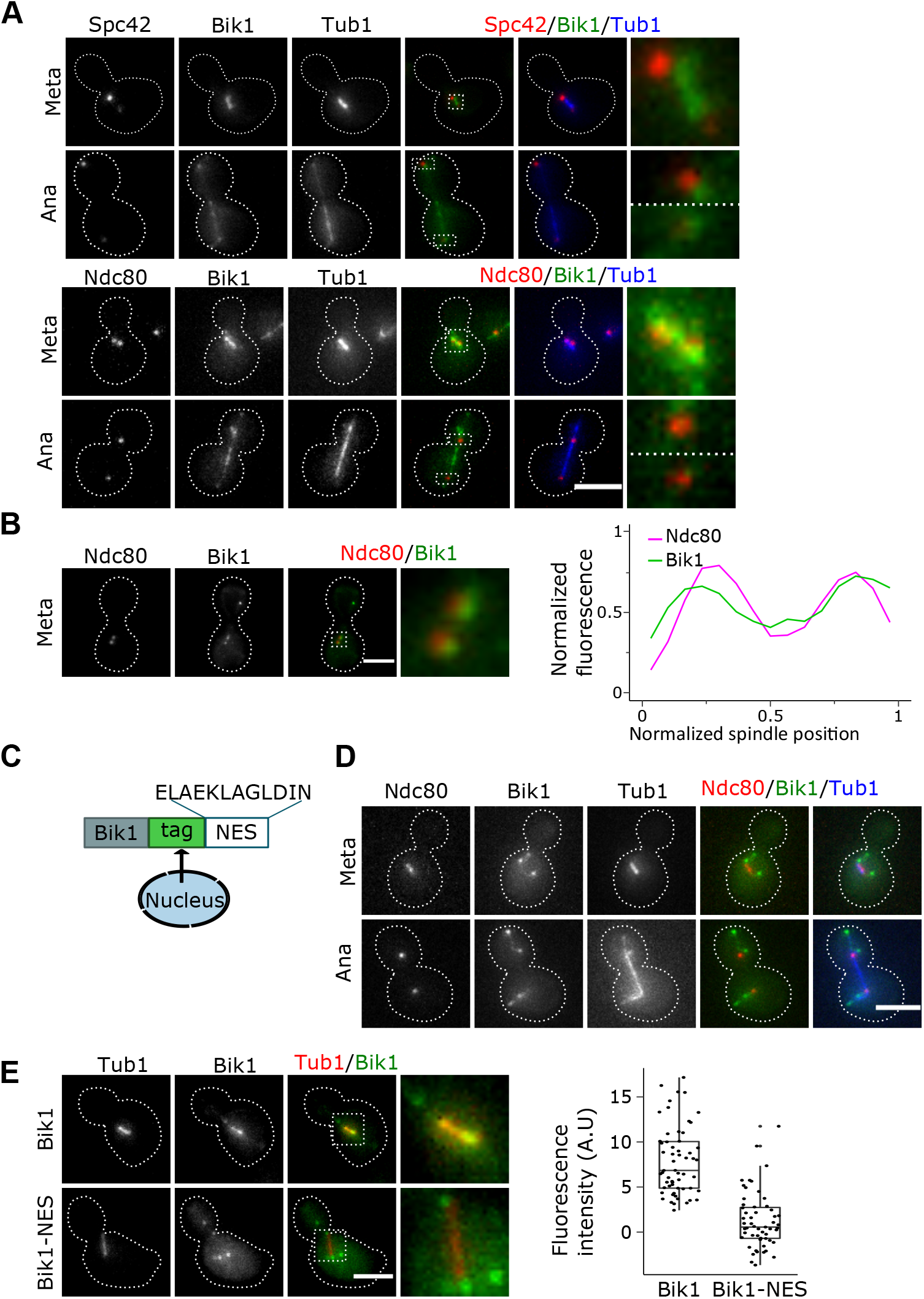
Bik1 localizes to kinetochores in a cell-cycle dependent manner. (A) Representative images of Bik1-sfGFP (green) localization in cycling cells expressing mTurquoise-Tub1 (blue) and Spc42-tagRFP-T (red, top panel), and Nc80-tagRFP-T (red, lower panel). Dashed boxes show area enlarged in the rightmost images. (B) Le panel: Representative images of Bik1-sfGFP (green) localization in pGal1-Cdc20 metaphase-arrested cells expressing Nc80-tagRFP-T. Right panel: Line scan analysis shows the distribution of Bik1-sfGFP and kinetochores (Ndc80-tagRFP-T) in metaphase arrested cells. For each cell, the fluorescence intensity was measured along a line drawn between the kinetochore foci and normalized relative to the maximum value, and the mean values are shown (n=26 cells). Dashed boxes show area enlarged in the rightmost images. (C) Cartoon showing the design of Bik1-NES. In short, a strong nuclear export signal (NES) was fused to Bik1-sfGFP or Bik1-5xFlag to promote nuclear export. (D) Representative images of Bik1-sfGFP-NES (green) localization in cycling cells expressing mTurquoise-Tub1 (blue) and Ndc80-tagRFPT (red). (E) Quanfication of Bik1-sfGFP or Bik1-sfGFP-NES localization in metaphase cells. Bik1-sfGFP and Bik1-sfGFP-NES cells expressing mRuby2-Tub1 were arrested in G1 with α-factor and subsequently released in fresh media. Cells were fixed every 10 minutes for 120 minutes. Left: Representative cells at metaphase. Time after release = 40 min (Bik1-sfGFP), and 50 min (Bik1-sfGFP-NES). Right: Boxplots of the integrated fluorescence intensity of Bik1-sfGFP and Bik1-sfGFP-NES at the spindle (mRuby2-Tub1) in metaphase cells (n=59 cells for Bik1-sfGFP and 55 for Bik1-sfGFP-NES). Dashed boxes show area enlarged in the rightmost images. All images shown are maximum intensity projections of Z-stacks (9 μm, 1 μm steps). Dashed lines represent the cell outlines based on bright field images. All scale bars, 4 μm.

To confirm the metaphase association with kinetochores, we used cells expressing the APC/C co-activator Cdc20 under the control of the galactose inducible *GAL1* promoter. Shifting the carbon source from galactose to glucose represses the expression of CDC20 and thereby arrests the cells at the metaphase-to-anaphase transition. Bik1-sfGFP was localized at kinetochore clusters in metaphase arrested cells (Figure 1B), confirming what we observed in cycling cells with short spindles. In summary, we conclude that Bik1 associates with kinetochores from G1 until metaphase, after which it instead is localized to interpolar microtubules.

### Nuclear Bik1 is needed for timely cell cycle progression

Bik1 in the cytoplasm localizes to astral microtubules and plays an important role in mitotic spindle positioning (Carvalho et al., 2004; Moore et al., 2006). To disrupt the kinetochore function of Bik1 while maintaining its cytoplasmic function, we fused a strong nuclear export consensus sequence (NES) (Kosugi et al., 2008) to the C-terminal end of Bik1-sfGFP or Bik1-5xFlag (collectively referred to as Bik1-NES, Figure 1C). We imaged Bik1-NES cells expressing mTurquoise-Tub1, and Nc80-tagRFPT and confirmed that the mutant indeed excludes Bik1 from the nucleus while maintaining its localization at astral microtubules (Figure 1D). To quantify the effectiveness of the NES, we followed the localization of Bik1 in Bik1-sfGFP and Bik1-sfGFP-NES cells expressing mTurquoise-Tub1. We synchronized the cells in G1 with α-factor, released them in fresh media, and fixed cells at 10-minute intervals from release. We then measured Bik1 intensity along the spindle in metaphase cells and found that, as expected, Bik1-NES cells have significantly less sfGFP signal at the spindle microtubules than Bik1 cells (Figure 1E).

We next assessed cell cycle progression in the absence of nuclear Bik1. We synchronized Bik1-sfGFP and Bik1-sfGFP-NES cells in G1 with α-factor and scored the number of cells in G1/S, metaphase, or anaphase according to the spindle length (mRuby2-Tub1) (Figure 2A). The strain expressing Bik1-sfGFP had a sharp peak of metaphase cells at 40 min and a high number of cells with long spindles 50-60 min after the release. Bik1-sfGFP-NES cells, by contrast, displayed a broader peak of metaphase cells that spans between 40-60 min post-release and a 10 min delay in the anaphase peak.

**Figure 2.**
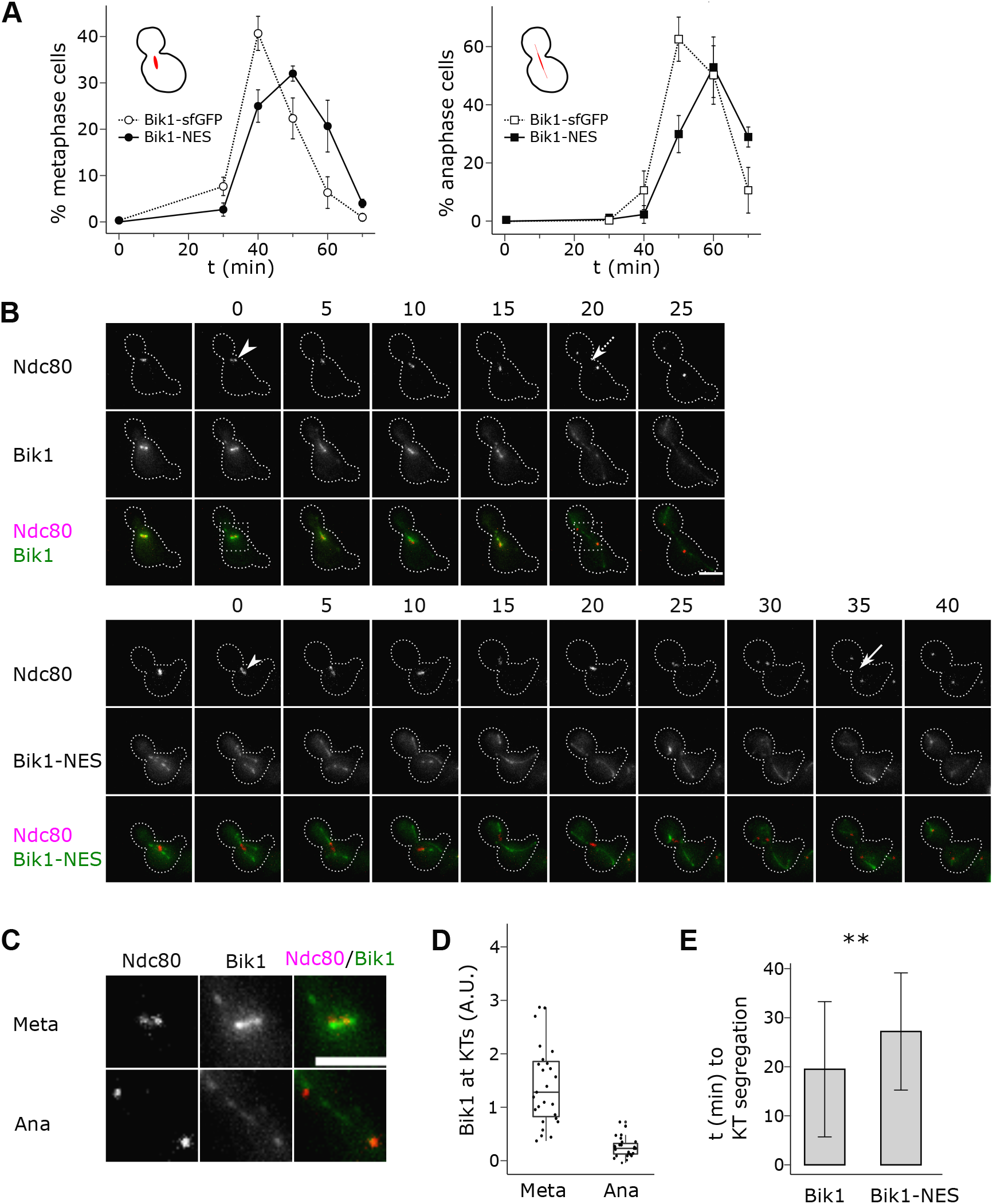
Bik1-NES delays spindle elongation at metaphase/anaphase transition. (A) Cells from (1E) were scored for cell cycle progression based on bud size and spindle length (mRuby-Tub1). Error bars represent mean ± sd (n=3 independent experiments, each experiment with 100 cells each mepoint for each strain (Bik1-sfGFP and Bik1-sfGFP-NES) analysed per experiment). (B-E) Time-lapse imaging of cells expressing Bik1-sfGFP or Bik1-sfGFP-NES (green) and Ndc80-tagRFP-T (red). Cells were arrested in G1 with α-factor and subsequently released in fresh medium under the microscope. Images were taken at 5 min intervals. (B) Representative me-lapse images. Arrowheads indicate the appearance of separated kinetochore clusters. Arrows indicate segregated kinetochores in mother/bud. The me (min) elapsed from appearance of separated kinetochore clusters is indicated by the numbers above the frames. Dashed boxes show area enlarged in (C). Images shown are maximum intensity projections of Z-stacks (9 μm, 1 μm steps). Scale bar, 4 μm. (C) Zoomed-in images from the dashed areas of (B), showing the kinetochores (Ndc80-tagRFP-T) at metaphase and anaphase. (D) Boxplots of the integrated Bik1-sfGFP fluorescence intensity at kinetochores (Ndc80-tagRFP-T) at metaphase (meta) and anaphase (ana) (n= 27 cells). (E) Quanfication of average me from the appearance of separated kinetochore clusters to segregated kinetochores in mother/bud in Bik1-sfGFP and Bik1-sfGFP-NES cells. We performed two independent experiments and pooled the data. Error bars represent mean ± sd (n= 59 cells for Bik1-sfGFP and 57 for Bik1-sfGFP-NES, p-value = 0.001691 using Welch Two Sample t-test (two-sided)).

We then examined Bik1-sfGFP and Bik1-sfGFP-NES cells after G1 synchronization with α-factor by time-lapse imaging (Figure 2B-E). In the Bik1-sfGFP strain, Bik1 was localized to the kinetochores from G1 through metaphase. When the spindle elongates in anaphase, Bik1 was no longer associated with the kinetochores and was found in patches along the interpolar microtubules instead (Figure 2B-C), as observed in fixed cells (Figure 1 A). The fluorescence intensity of Bik1-sfGFP at kinetochores was highest at metaphase when kinetochores cluster and reduced to near background levels in anaphase (Figure 2D). In agreement with the cell-cycle analysis from fixed cells, we observed a slight delay in cell cycle progression in Bik1-NES cells (Figure 2B, arrows). Our quantifications showed that the time from the appearance of two kinetochore clusters to their segregation into mother and daughter cells is approximately 10 min longer in Bik1-NES cells (Figure 2E). Interestingly, we occasionally observed all kinetochores of short spindles transferred to the daughter cell in the NES mutant strain before elongation occurred (Figure 2 B, 15 min and 25-30 min). We interpret this as a secondary effect of the prolonged metaphase of the mutant.

### Cells lacking nuclear Bik1 display aberrant kinetochore positioning

Because Bik1 localizes near the kinetochores in metaphase, we hypothesized that the prolonged metaphase of Bik1-NES cells might be due to a dysregulation in kinetochore clustering. To test this hypothesis, we analyzed the distribution of the kinetochore component Ndc80-sfGFP relative to the SPBs (Spc42-tagRFPT) in cells expressing either Bik1-5xFlag or Bik1-5xFlag-NES arrested in metaphase (PGal1-Cdc20) (Figure 3). While Bik1-5xFlag cells showed typical Ndc80 clusters, we observed cells with long stretches of Ndc80 in the Bik1-5xFlag-NES mutant (Figures 3A-B). The unclustered kinetochore phenotype was observed in 60% of the Bik1-NES metaphase cells (Figure 3C). Line scan analysis of the Ndc80-sfGFP fluorescence relative to the SPBs showed that Bik1-NES cells have a flattened distribution of Ndc80 with kinetochores mispositioned towards the spindle equator (Figure 3D).

**Figure 3.**
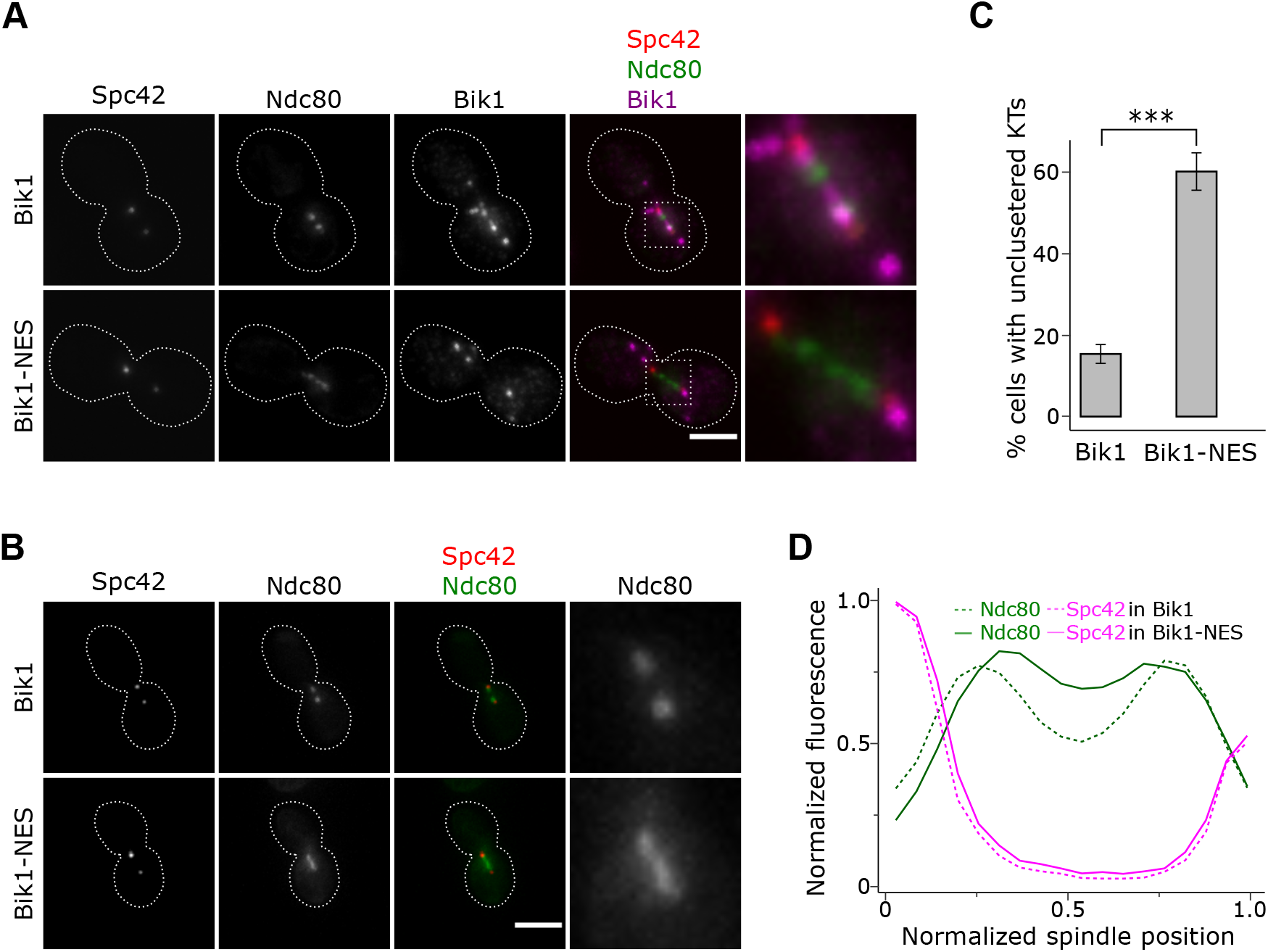
Nuclear Bik1 is needed for kinetochore clustering at metaphase. (A) Representative immunofluorescence staining for Bik1-5xFlag and Bik1-5xFlag-NES (magenta) in metaphase arrested pGal1-Cdc20 cells expressing Spc42-sfGFP (red) and Ndc80-TagRFP-T (green). Images shown are maximum intensity projections of Z-stacks (4.95 μm, 0.3 μm steps). Dashed boxes show area enlarged in the rightmost images. Scale bar, 4 μm. (B) Representative images of Bik1-5xFlag and Bik1-5xFlag-NES arrested as in (A), including zoomed-in images of the kinetochores (Ndc80-sfGFP). Images shown are maximum intensity projections of Z-stacks (8.7 μm, 0.3 μm steps). Scale bar, 4 μm. (C) Quanfication of kinetochore (Ndc80-tagRFPT) unclustered phenotype in Bik1-5xFlag and Bik-5xFlag-NES cells arrested in metaphase as shown in (B). Error bars represent mean ± sd (n=3 independent experiments, with 100 cells for each strain (Bik1-5xFlag and Bik1-5xFlag-NES). p=0.0006884 using Welch Two Sample t-test (two-sided). (D) Line scan analysis shows the distribution of kinetochores (Ndc80sfGFP) and spindle pole bodies (SPBs, Spc42-tagRFP-T) in metaphase arrested cells. For each cell, the fluorescence intensity was measured along lines drawn between the SPBs (Spc42-TagRFP-T) and normalized relative to the maximum value. Mean values are shown (n=100 cells for each strain (Bik1-5xFlag and Bik1-5xFlag-NES).

Bik1 has previously been suggested to contribute to the kinetochore-microtubule attachment. Bik1 is needed to nucleate microtubules that capture kinetochores after centromere re-activation (Tanaka et al., 2005). Furthermore, a binding partner of Bik1, the +TIP Bim1, was recently shown to promote *in vitro* oligomerization of Dam1 complexes– a major component of the yeast outer kinetochore (Dudziak et al., 2021). Therefore, Bik1’s function in kinetochore end-on attachment could contribute to the cell cycle delay we observe. Yet, we did not detect unattached kinetochores in the Bik1-NES mutant. Instead, the kinetochores are aligned in the spindle but appear unclustered during metaphase.

Previous studies have shown that the kinesins Cin8 and to a lesser extent Kip1 and Kip3 are required for yeast chromosome congression (Gardner et al., 2008; Tytell & Sorger, 2006; Wargacki et al., 2010). However, the deletion of each kinesin results in slightly different types of clustering defects. In the cin8Δ mutant, the kinetochores are positioned closer to the spindle equator (Gardner et al., 2008; Tytell & Sorger, 2006; Wargacki et al., 2010). The kip1Δ mutant has a similar, but weaker, phenotype (Gardner et al., 2008; Tytell & Sorger, 2006). Conversely, Ndc80 clusters in kip3Δ cells appear closer towards the spindle poles (Wargacki et al., 2010). We found that the Ndc80 foci in the Bik1-NES mutant appear shifted towards the spindle equator (Figure 3D), as has been reported for cin8Δ cells.

### Bik1 interacts with Cin8 at the spindle

To investigate Bik1’s association with kinesins, we performed single-step affinity purification of Bik1-Flag in metaphase arrested cells, followed by mass spectrometry analysis. Bik1-Flag co-purified with the yeast kinesins Kip2, Cin8, Kip1, Kip3 and Kar3 (Table 1). Notably, the kinesin Cin8 appeared with a similar coverage as the known interactor Kip2 (Supplementary data S1).

The defect of Bik1-NES cells in kinetochore clustering shown in Figure 3A phenocopies CIN8 deletion (Gardner et al., 2008; Tytell & Sorger, 2006; Wargacki et al., 2010). Therefore, we decided to study the Bik1-Cin8 interaction in more detail. We investigated whether Cin8 and Bik1 interact *in vivo* using proximity-dependent biotin identification (TurboID), a proximity labelling assay that can detect transient or weak protein-protein interactions (Roux et al., 2012). We fused TurboID, a mutated version of a bacterial biotin ligase BirA (Branon et al., 2018; Larochelle et al., 2019), to the C-terminus of Bik1 (Figure 4A). We then Flag-tagged Cin8 or Kip2, a kinesin that interacts with Bik1 at astral microtubules (Carvalho et al., 2004). Both Cin8 and Kip2 were recovered after streptavidin pulldown when expressed in Bik1-TurboID cells, indicating that Bik1 and Cin8 are in proximity *in vivo* (Figure 4B).

**Figure 4.**
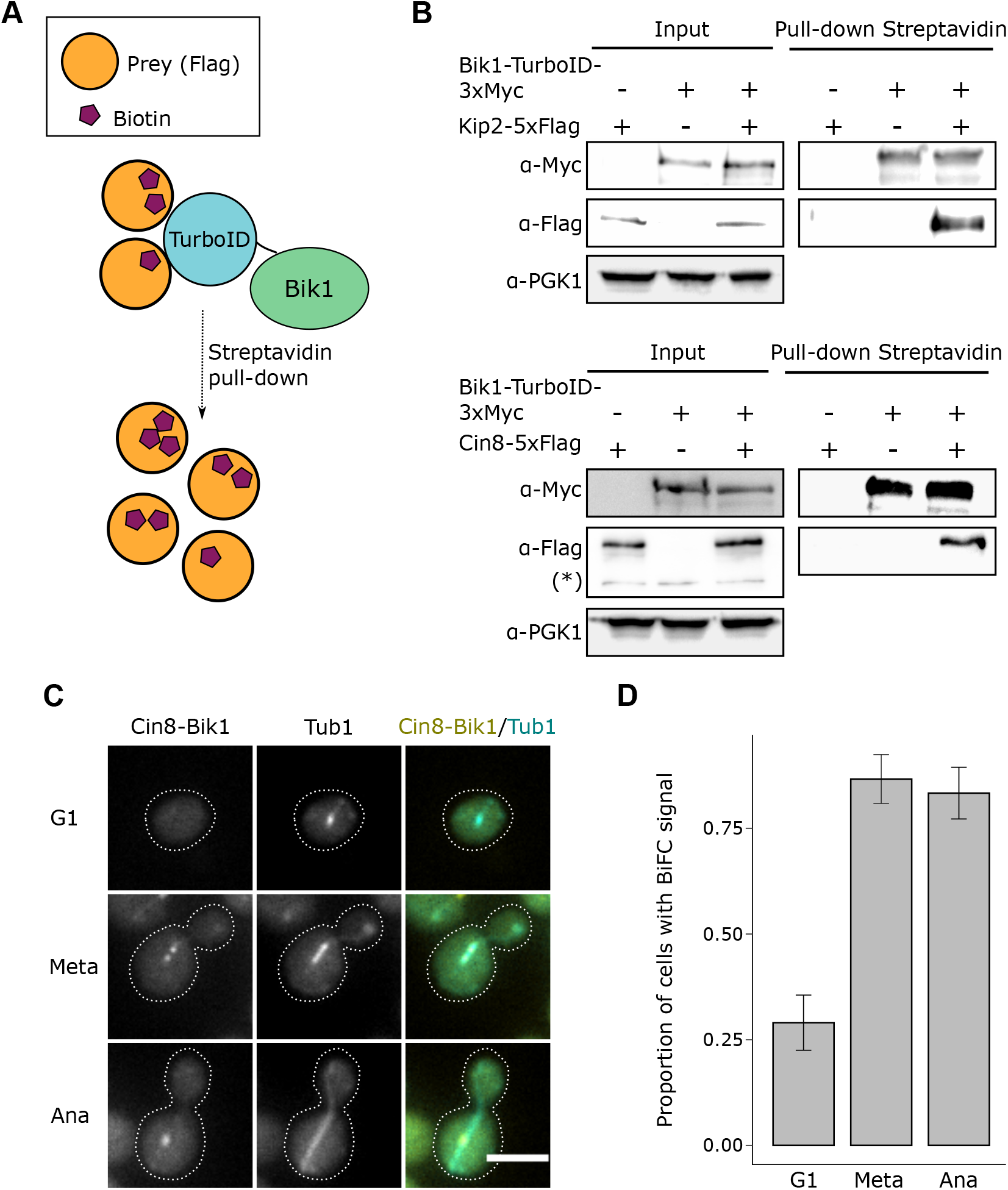
Bik1 interacts with Cin8 at metaphase. (A-B) Idenfication of Cin8 as a Bik1 interaction partner using TurboID. Cartoon outlining the principle of the TurboID assay. Briefly, a biotin ligase was fused to endogenous Bik1 (Bik1-TurboID) in cells expressing the protein of interest tagged with Flag (Prey). Biotinylated substrates were purified using streptavidin beads, and inputs and pull-down lysates were analyzed by immunoblot for the indicated proteins. (B) Western blot analysis of total cell extracts (input) and streptavidin pull-downs prepared from Bik1-TurboID-3xmyc cells expressing Kip2-5xFlag or Cin8-5xFlag as prey, and control strains, using anti-Myc (top) and anti-Flag (bottom) antibodies. (*) unspecific band. (C-D) Bimolecular fluorescence complementation (BiFC) analysis of Bik1 and Cin8 interaction. Representative images of cycling cells expressing Bik1-VN and Cin8-VC in G1, metaphase and anaphase. Images shown are single focal planes. (D) Proportion of cells in (C) with BIFC signal based on Venus fluorescence. Error bars represent mean ± sd (n= 3 independent experiments, with 50 cells analyzed in each cell cycle phase (G1, metaphase and anaphase) per experiment.

We next looked at the interaction between Bik1 and Cin8 in individual cells by bimolecular fluorescence complementation (BiFC) (Sung & Huh, 2007). Briefly, an N-terminal fragment of the yellow fluorescent protein Venus was fused to the C-terminus of Bik1, and a complimentary C-terminal Venus fragment was fused to the C-terminus of Cin8. We detected BiFC signal at the spindle, predominantly in metaphase and anaphase cells (Figure 4 C-D). Interestingly, the BiFC signal in metaphase cells appeared as 1-2 dots at the spindle, while the signal in anaphase cells appeared as a single dot around the spindle midzone (Figure 4 C).

We then investigated the dependence of Cin8 and Bik1 to localize to the spindle. In the Bik1-NES mutant, Cin8 still forms two foci between SPBs and kinetochores (Figure 5A and B), even in cells with unclustered kinetochores. This result was surprising as Cin8 has been shown to bind kinetochores directly (Suzuki et al., 2018). Linescan analysis of Cin8-sfGFP localization in relation to SPBs showed that Cin8 localization is unaltered in the absence of nuclear Bik1 (Figure 5B). Deletion of CIN8, on the other hand, resulted in a significantly reduced Bik1 intensity at the spindle (Figure 5C-D). Overall, these data suggest that a portion of nuclear Bik1 associates with and is dependent on Cin8 to bind the spindle.

**Figure 5.**
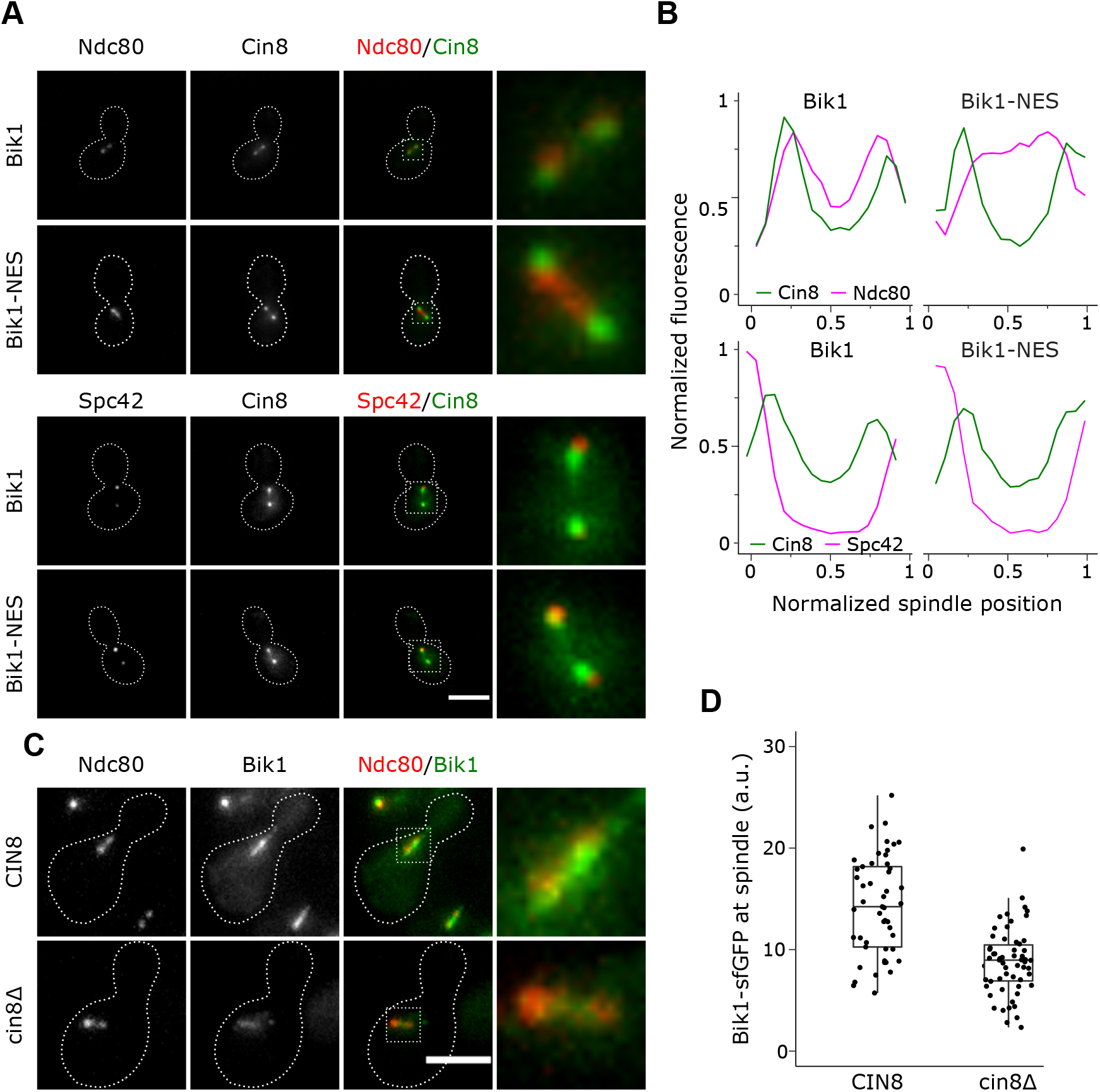
Dependence of Cin8 and Bik1 to localize to the spindle. (A-B) Cin8 localizes at the spindle in Bik1-NES cells but does not co-localize with unclustered kinetochores. (A) Cin8 localization at the metaphase spindle does not require nuclear Bik1. Representative images of pGal1-Cdc20 cells arrested in metaphase expressing Bik1-5xFlag or Bik1-5xFlag-NES, Cin8-sfGFP, and the markers Ndc80-TagRFP-T (top panel) or Spc42-TagRFP-T (lower panel). Images shown are maximum intensity projections of Z-stacks (8.7 μm, 0.3 μm steps). Dashed boxes show area enlarged in the rightmost images. Scale bar, 4 μm. (B) Line scan analysis of Cin8 distribution (Cin8-sfGFP) in metaphase arrested cells). Top panel: Cin8 distribution relative to the kinetochores (Ndc80-tagRFP-T). For each cell, the fluorescence intensities of Cin8-sfGFP and Ndc80-tagRFPT was measured along lines drawn between the SPBs and normalized relative to the maximum values, mean values are shown (n= 25 cells for BIK1 and 25 for Bik1-5xFlag-NES). Low panel: Cin8 distribution relative to the SPBs (Spc42-tagRFP-T). For each cell, the fluorescence intensity of Cin8-sfGFP and Spc42-tagRFPT was measured along a line drawn between the SPBs and normalized relative to the maximum value. Mean values are shown (n= 25 cells for Bik1-5xFlag and 26 for Bik1-5xFlag-NES). (C) Cin8 is needed for Bik1-sfGFP localization at kinetochores. CIN8 and cin8Δ cells expressing Bik1-sfGFP and Ndc80-TagRFP-T were arrested in G1 using α-factor, released in fresh media at 25°C and fixed every 10 minutes for 130 minutes. Left: Representative images of cells in metaphase (CIN8, 60 min after release; cin8Δ, 110 min after release). Images shown are maximum intensity projections of Z-stacks (6.6 μm, 0.6 μm steps). Dashed boxes show area enlarged in the rightmost images. Scale bar, 4 μm. (D) Boxplots of the integrated fluorescence intensity of Bik1-sfGFP at the spindle (based on Ndc80-TagRFP-T localization) in metaphase cells (n=100 cells for each strain (CIN8 and cin8 Δ).

The reduced Bik1 recruitment to the spindle and kinetochores explains why Bik1 cannot compensate for the lack of Cin8, which is known to cause severe defects in assembling the mitotic spindle (Hoyt et al., 1992; Kotwaliwale et al., 2007). A previous study has shown that Cin8 mediates chromosome congression by depolymerizing kinetochore microtubules in a length-dependent manner (Gardner et al., 2008). Gardner and coworkers proposed that Cin8 binds randomly to kinetochore microtubules and walks towards the plus-end to promote microtubule plus-end disassembly directly. However, in Bik1-NES cells, Cin8 is insufficient to regulate chromosome congression. Moreover, Cin8 does not localize to the unclustered kinetochores in the Bik1-NES mutant. Taken together, these observations suggest that Bik1 is a binding partner of Cin8 that contributes to the disassembly of kinetochore microtubules to achieve chromosome congression at metaphase. There are at least two other examples where plus-end directed kinesin motors interact with Bik1. Kip2 transports Bik1 to astral microtubule plus ends to ensure proper spindle alignment (Carvalho et al., 2004; Caudron et al., 2008). Furthermore, it has been suggested that Bik1 and Kar3 form a complex at the junction of oppositely oriented MTs during mating to induce microtubule depolymerization for nuclear congression to occur (Molk et al., 2006).

We propose that Bik1 increases the processivity of Cin8. Bik1 has been suggested to be a processivity factor for the kinesin Kip2 at astral microtubules (Hibbel et al., 2015). Likewise, Bik1 might be needed for Cin8 to reach the plus-ends of long kinetochore microtubules that extend beyond bundled kinetochore microtubules. Cin8 is a bidirectional kinesin and *in vitro* studies have shown it preferentially moves in a minus-end directed manner as single motors (Gerson-Gurwitz et al., 2011; Pandey et al., 2021; Roostalu et al., 2011). Directional switching occurs when Cin8 motors are clustered (Pandey et al., 2021; Roostalu et al., 2011), and other kinesin-5 motors switch directionality in response to steric crowding on MTs (Britto et al., 2016). Loss of nuclear Bik1 could therefore lead to ineffective plus-end directed movement of Cin8 on long kinetochore microtubules and failure to cluster kinetochores at metaphase. This is supported by the observation that Cin8 is not localized along rod-like distribution of Ndc80 in the Bik1-NES mutant.

## Materials and Methods

### Yeast strains and plasmids

All *S. cerevisiae* strains used in this study are derivatives of BY4741 and are listed in Table S1. The epitope tagged alleles (sfGFP and tagRFPT), TurboID-3xmyc and BiFC were constructed at the endogenous loci by standard PCR-based integration as described in (Longtine et al., 1998) and confirmed by PCR, sequencing and microscopy. The plasmids used to generate the strains are listed in Table S2. Strains with fluorescently tagged Tub1 were constructed as described in (Markus et al., 2015). Briefly, the listed plasmids were linearized using BsaBI (New England Biolabs, R0537S) and integrated at the endogenous TUB1 locus by homologous recombination.

### Growth conditions

Cells were grown at 30°C in YEP (1% yeast extract, 2% bacto-peptone) medium supplemented with 2% glucose (YEPD) unless otherwise stated. For G1 arrest, overnight cultures were diluted to OD_600_ = 0.1 in fresh media, grown for 2.5 hours and arrested by adding 3 μg/ml α-factor (Sigma, custom peptide WHWLQLKPGQPMY) every 60 min for 120 min. Cells were released by washing away α-factor with 1x culture volume of YEPD and resuspending in 1x culture volume of YEPD, and further grown at 30°C. For metaphase arrest, PGal1-Cdc20 cells were grown in YEP supplemented with 2% raffinose and 2% galactose, and switched to YEPD for 120 min.

### Fluorescence imaging

Cells were fixed in 4% formaldehyde on ice for 60 minutes, washed with PBS and stored in PBS at 4°C until imaging. Cells were imaged in concanavalin A-coated 96 well glass bottom plates. Images were acquired using a Nikon Ti microscope, equipped with a Zyla 4.2+ sCMOS camera and a 60x water immersion objective (NA 1.2). Z-stack images were captured for fixed cells, as described in the figure legends. Images for BiFC and immunofluorescence were acquired using a Nikon Ti2 microscope, equipped with a Prime BSI sCMOS camera and a 60x water immersion objective (NA 1.2).

### Live-cell imaging

Cells were arrested in G1 with as α-factor and subsequently released in low-fluorescence media (LFM) containing 1.7 g/l yeast nitrogen base without ammonium sulfate, amino acids, folic acid and riboflavin (MP-Biomedicals, 4030-512), and 1 g/l monosodium glutamate, supplemented with 2% glucose and amino acids. Cells were left to settle on the bottom of concanavalin A-coated 96 well glass bottom plates for 20-30 minutes inside the microscope incubator at 30°C before being imaged. Images were captured every 5 min for 100-120 min.

### Image analysis

Fluorescence intensity measurements were made from maximum intensity projections of image z-stacks using FiJi (Schindelin et al., 2012). Integrated intensities were measured in a region of interest drawn around spindles/kinetochores. Background correction was done by multiplying the mean fluorescence of regions outside of the nucleus with the measured region’s area and subtracting from the measured integrated intensity. Linescan analysis was performed by drawing a line between the centers of opposing SPB foci and measuring the fluorescence intensities along the line. Background correction was done by subtracting the mean fluorescence measured at the cytoplasm. Binned means were calculated for each strain. Cell cycle analyses were performed by classifying the cells according to the spindle length, based on the Tub1-tagRFPT marker. (G1/S) spindles <1 μm; metaphase (1-2 μm), and anaphase (>2 μm).

### Immunofluorescence

PGal1-Cdc20 cells were arrested in metaphase and fixed by adding formaldehyde to 3.7% (vol/vol) and incubating while shaking at 30°C for 30 min. The fixed cells were washed once with PBS and then twice with SP buffer (0.1 M KPO4, 1.2 M sorbitol, pH 7.5). The cells were resuspended in 500 μl SP buffer with 0.1% β-mercaptoethanol and 10 μl of 20 mg/ml Zymoylase-20T (Seikagaku). The cells were incubated with soft rotation for 20 min at room temperature (RT). The resulting spheroplasts were then washed twice in SP buffer and permeabilized in blocking solution (3% BSA 0.1% Triton X-100 in PBS) for 30 min at RT. The spheroplasts were immunostained in suspension. First, cells were incubated with 1:1000 mouse monoclonal anti-FLAG M2 antibody (Sigma, F1804) in blocking buffer for 1 hour at RT. Cells were washed three times in PBS and then incubated with 1:100 donkey anti-mouse Alexa Fluor 647 antibody (ThermoFisher, A-31571) for 45 min at RT. Cells were washed three times in PBS, seed in a glass-bottom imaging plate coated with Concanavalin A, and imaged.

### Sample preparation for mass spectrometry analysis

Bik1-5xFlag PGal1-Cdc20 cells were arrested in metaphase. Briefly, cells were grown overnight in 50 ml YEP with 2% galactose and 2% raffinose. Cells were diluted in 2 L YEP with 2% and 2% raffinose and grown at 30°C to OD_600_ = 1. Then, cells were spun down and resuspended in 2 L YEPD with 2% glucose and arrested at 30°C for 2 h. Cells were spun down at 6000 rom at 4°C, washed with cold dH_2_O, and spun down at 6000 rpm at 4°C. The cell pellet was diluted in a few drops of lysis buffer (50 mM Tris, pH 8.0, 150 mM NaCl, 5% glycerol, 0.1% Triton X-100). The cell suspension was snap frozen by pipetting the drops directly into liquid nitrogen. The frozen pellets were then lysed using a Freezer/Mill (SPEX™ sample prep freezer/mill), using 10 rounds consisting of 2 min grinding at 14 rpm, 2 min cooling). The yeast pellets were diluted in buffer EB1 (40 mM sodium hepes, 300 mM NaCl, 0.5% Triton X-100, 2 mM MgCl_2,_ 5% glycerol and protease and phosphatase inhibitors (1mM PMSF, 1mM leupeptin, 1mM peptasin A, 10mM NaF, 1mM sodium pyrophosphatase, 1mm sodium orthovanadate), and clarified by centrifugation at 10 000 g for 10 min at 4 °C. The cleared lysate was incubated for 30 min with magnetic Dynabeads (Thermo Fisher Scientific) coupled with anti-FLAG M2 antibody (Sigma-Aldrich). The Dynabeads were washed three times with buffer A (25 mM sodium Hepes, 2 mM MgCl_2_, 0.1% EDTA, 0.1% Triton X-100, 5% glycerol, 150 mM NaCl, protease and phosphatase inhibitors (1 mM PMSF, 1 mM leupeptin, 1 mM peptasin A, 10mM NaF, 1 mM sodium pyrophosphatase, 1 mm sodium orthovanadate) and four times with buffer B (25 mM Hepes, pH 8.0, and 150 mM KCl).

### Protein on-beads digestion

Proteins bound to the Dynabeads were resuspended and washed with 500 μL of 50 mM ammonium bicarbonate (AmBic), pH 8.0, twice. After incubation at room temperature for 15 min, 50 μL of 50 mM AmBic was added. Protein disulfide bonds were reduced with 5 μL of 200 mM dithiothreitol at 37°C for 45 min and alkylated with 5 μL of 600 mM iodoacetamide at room temperature in dark for 30 min. Tryptic digestion was started by adding 1 μg of trypsin and incubation at 37°C overnight, followed by adding another 500 ng of trypsin and incubation at 30°C for 3 h. The digestion was stopped with 5 μL concentrated formic acid. The samples were then cleaned on a C18 StageTip (Thermo Scientific) and dried using a speedvac (MiVac, Thermo Scientific).

### PRLC-MS/MS analysis

Chromatographic separations of peptides were performed on a 50 cm long C18 EASY-spray column connected to an Easy nanoLC-1000 UPLC system (Thermo Fisher Scientific). The gradient was 4-25% of solvent B (98% acetonitrile, 0.1% formic acid) in 50 min and 25-40%B in 10 min at a flow rate of 300 nl/min. An Orbitrap Fusion mass spectrometer (Thermo Scientific) analyzed the eluted peptides that were ionized in a nano-Easy electrospray source. The survey mass spectrum was acquired at the resolution of 120,000 in the range of *m/z* 350-1800. MS/MS data of precursors were obtained with 30% normalized collision energy by higher-energy collisional dissociation (HCD) for ions with charge *z*=2-7 at a resolution of 30,000 in 3 s cycle time.

### Data analysis

Mascot v 2.5.1 search engine (MatrixScience Ltd., UK) was used for protein identification with the following search parameters: up to two missed cleavage sites for trypsin, peptide mass tolerance 10 ppm, 0.05 Da for the HCD fragment ions. Carbamidomethylation of cysteine was specified as a fixed modification, whereas oxidation of methionine, deamidation of asparagine and glutamine were defined as variable modifications.

### Turbo-ID assay

Overnight cultures were diluted to OD_600_ = 0.2 in YEPD medium supplemented with 50 μM biotin and grown at 30°C until they reached OD_600_ = 0.6 (4-5 h). Cells were pelleted and resuspended in RIPA buffer (50 mM Tris-HCl, 150 mM NaCl, 1 mM EDTA, 0.1% SDS, 1% Triton X-100, 1.5 mM MgCl2, 0.4% sodium deoxycholate, 1 mM DTT, 1 mM PMSF, 1 mM leupeptin and1 mM pepstatin A). The cells were lysed using acid-washed glass beads and a bead-beater at 4°. To shear the DNA and RNA, the cell lysate was sonicated, and then clarified by centrifugation. Clarified lysates were incubated with 100 μl of Streptavidin-Sepharose beads (GE Healthcare; 17-5113-01) for 3 h at 40°C. The beads were then washed once with wash buffer (50mM Tris-HCl, 2% SDS) and three times with RIPA buffer. The purified proteins were eluted with 4 x Laemmli buffer containing 25 μM biotin and 10% β-mercaptoethanol, for 15 min at 95°C. The whole cell extract or pull-down samples were separated by SDS-PAGE, and the proteins were transferred to a PVDF membrane (BioRad) followed by and standard immunoblotting. Primary (monoclonal anti-FLAG M2 antibody (Sigma-Aldrich, F3165, diluted 1:5000); c-Myc rabbit polyclonal antibody (Santa Cruz Biotechnology; sc-789, diluted 1:5000); anti-Pgk1 mouse monoclonal antibody (Invitrogen, 45925, diluted 1:5000), and secondary anti-mouse IgG peroxidase antibody, (Sigma, A9044, diluted 1:50000) were used. HRP conjugated secondary antibodies were detected with Femto ECL (ThermoFisher Scientific).

## ABBREVIATIONS

(ATP): Adenosine triphosphate
(NES): nuclear export signal
(SPB): spindle pole body
(+TIPs): microtubule plus-end tracking proteins
(sfGFP): superfolder GFP
(BiFC): bimolecular fluorescence complementation

## Acknowledgements

We thank C. Björkegren (Karolinska Institutet, Sweden) for reagents, and C. Boone and Z. Li for help with construction of deletion mutants and critical reading of this manuscript. Microscopy was performed at the Live Cell Imaging Core facility/Nikon Center of Excellence, at the Karolinska Institutet, supported by grants from the Swedish Research Council, KI infrastructure, and Centre for Innovative Medicine. The BioImage Informatics Facility at SciLifeLab, National Microscopy Infrastructure (VR-RFI 2019-00217), and recipient of the Chan-Zuckerberg Initiative, assisted in the image analysis. Protein identification and quantification were carried out by the Proteomics Biomedicum core facility, Karolinska Institutet. This work was supported by grants from the Swedish Council (VR-NT 2017-04536), and STINT (Mobility Grant for Internationalization, MG2019-8484) to V.M.B. The authors declare no competing financial interests.

---

**Table S1.**
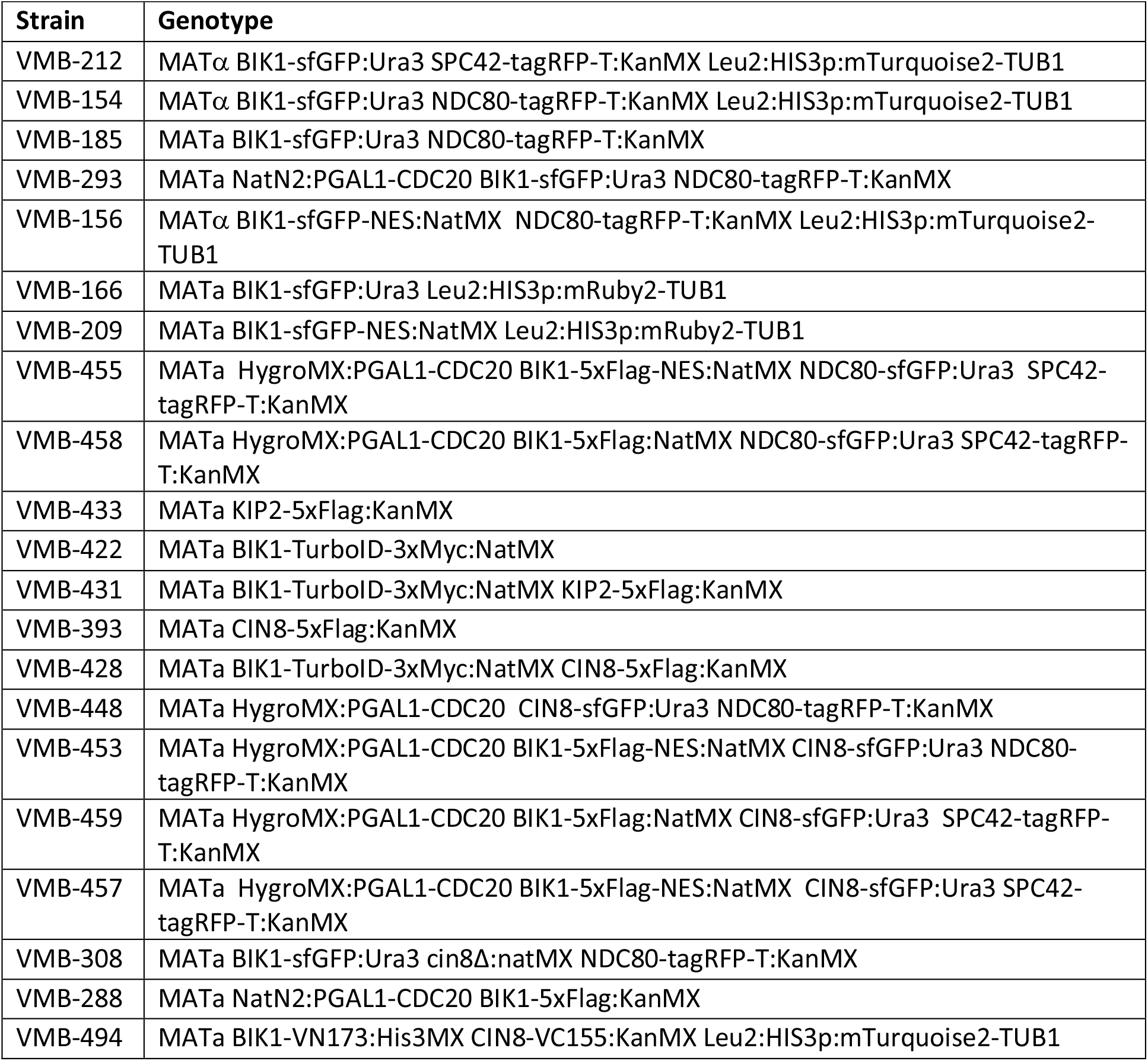
Strains used in this study. All strains are derived from BY4741

**Table S2.** Plasmids used in this study

| Plasmid | Relevant insert | Source |
| --- | --- | --- |
| p44873 | pFA6a-link-yosuperfolderGFP-CaURA3 | Gift from Wendell Lim & Kurt Thorn (Addgene: 44873) PMID: 23844123 |
| p44906 | pFA6a-link-yoTagRFP-T-Kan | Gift from Wendell Lim & Kurt Thorn (Addgene: 44906) PMID: 23844123 |
| p50641 | pHIS3p:mTurquoise2-Tub1+3'UTR::LEU2 | Gift from Wei-Lih Lee (Addgene: 50641) PMID: 25711127 |
| p50645 | pHIS3p:mRuby2-Tub1+3'UTR::LEU2 | Gift from Wei-Lih Lee (Addgene: 50645) PMID: 25711127 |
| pYM-N23 | ScGAL1 promoter::pAgTEF-nat-tScADH1 | PCR-Toolbox Knop (Euroscarf) MID: 15334558 |
| p15983 | pFA6a-5xFlag-KanMX6 | Gift from Eishi Noguchi (Addgene: 15983) PMID: 18729046 |
|  | pFA6a-5xFlag-NatMX6 | This study (subcloned from p15983) |
| pFB1434 | pFA6a-Turbo-ID-3Myc-NatMX6 | Gift from François Bachand (Addgene: 126050) PMID: 31064814 |
| P30555 | pFA6a-VN-His3MX6 | Gift from Sung MK & Huh WK. (Euroscarf) PMID: 17534848 |
| P30560 | pFA6a-VC-kanMX6 | Gift from Sung MK & Huh WK. (Euroscarf) PMID: 17534848 |

